# Choice of Contact Points Modulates Sensorimotor Cortical Interactions for Dexterous Manipulation

**DOI:** 10.1101/621466

**Authors:** Pranav J. Parikh, Justin M. Fine, Marco Santello

## Abstract

Humans are unique in their ability to perform dexterous object manipulation in a wide variety of scenarios. However, previous work has used a grasping context that predominantly elicits memory-based control of digit forces by constraining where the object should be grasped. For this ‘constrained’ grasping context, primary motor cortex (M1) is involved in storage and retrieval of digit forces used in previous manipulations. In contrast, when choice of digit contact points is allowed (‘unconstrained’ grasping), behavioral studies revealed that forces are adjusted, on a trial-to-trial basis, as a function of digit position. This suggests a role of online feedback that detects digit position, rather than memory, for force control. However, despite the ubiquitous nature of unconstrained hand-object interactions in activities of daily living, the underlying neural mechanisms are unknown. Using non-invasive brain stimulation and electroencephalography, we found the role of M1 to be sensitive to grasping condition. While confirming the role of M1 in storing and retrieving learned digit forces and position in ‘constrained’ grasping, we also found that M1 is involved in modulating digit forces to digit position in unconstrained grasping. Furthermore, we found that digit force modulation to position relies on sensorimotor integration mediated by primary sensory cortex (S1) and M1. This finding supports the notion of a greater contribution of somatosensory feedback of digit position in unconstrained grasping. We conclude that the relative contribution of memory and online feedback based on whether contact points are constrained or unconstrained modulates sensorimotor cortical interactions for dexterous manipulation.

## INTRODUCTION

Dexterous object manipulation is a hallmark of human evolution (1–4). Co-adaptation of anatomical features and sensorimotor control mechanisms (5) have made dexterous manipulation a versatile means of interacting with the environment, while inspiring (6) and challenging (7) efforts to build dexterous robotic and prosthetic hands. The ability to skillfully use our hands depends on cortical mechanisms supporting several sensorimotor processes (4, 8–10), including integration of sensorimotor memory of previous hand-object interactions with online sensory feedback (11, 12). Although the role of motor and parietal cortices in this sophisticated interplay has been extensively studied (13–17), this work has drawn an incomplete picture of these cortical mechanisms. Research over the past three decades has focused on the control of digit forces through a paradigm based on grasping objects at visually-cued contacts (*constrained* grasping) (12, 18). These studies have shown that subjects use the same digit forces over consecutive trials by relying on a sensorimotor memory (19–22). Upon lifting the object, online sensory feedback is used to assess the accuracy of the force plan and update the sensorimotor memory of digit forces for future manipulations if an error occurs, e.g., object slip or tilt (11, 12, 19, 20, 22). The ‘constrained grasping’ paradigm, while providing significant insights into neural control of object manipulation, has neglected a critical component of sensorimotor control that is fundamental to natural hand-object interactions: choice of contact points.

When individuals can choose where to grasp an object (*unconstrained* grasping) – as it happens in many activities of daily living –, the central nervous system is presented with unique challenges: as there are no visual cues constraining where to grasp an object, unconstrained grasping is characterized by greater trial-to-trial variability of digit position than constrained grasping; this occurs even after the object dynamics have been fully learned (23–28). If control of digit forces in unconstrained grasping relied predominantly on sensorimotor memory, the same forces would be applied on each trial regardless of contact point. This behavior would lead to task failure. Remarkably, skilled manipulation can still be accurately performed because participants modulate digit forces as a function of digit position on a trial-to-trial basis (25–27). This evidence suggests that individuals do not rely primarily on memory of digit forces in unconstrained grasping. We have proposed that the predominant mechanism involves online feedback of digit position to change the force distribution every time an object is grasped at novel contact points (25, 26).

The ability to modulate digit forces to position raises the question as to whether adding choice of digit placement to manipulation would elicit distinct interactions among cortical grasp regions. Allowing choice of contact points has revealed differences in brain activation (29) and corticospinal excitability (30). However, the causal role of primary motor and somatosensory cortices (M1 and S1, respectively) for the control of digit forces and position remains to be established. The present work was designed to address this issue by combining brain stimulation, electroencephalography, and a dexterous manipulation paradigm we have validated in our previous studies.

Based on the above behavioral evidence (25–27) we hypothesized that, in unconstrained grasping, M1 and S1 are both involved in digit force-to-position modulation, each area being responsible for distinct functions: S1 would relay somatosensory feedback about digit position to M1, while M1 would process this feedback to modulate digit forces. Therefore, we predicted that a virtual lesion to M1 or S1 in unconstrained grasping should interfere with digit force-to-position modulation. In contrast, in constrained grasping a virtual lesion to M1 should only impair retrieval of digit forces used in previous manipulations. This hypothesis is based on evidence implicating M1 with building, storing, and retrieving sensorimotor memories of grasp forces in constrained grasping (15, 17). Lastly, based on previous work (31) we expected a virtual lesion to S1 to have no effect on digit forces.

Our experiments uncovered important functional differences in brain dynamics over M1 and S1 between the two grasp contexts. We first demonstrate that effective connectivity from S1 to M1 is significantly stronger in unconstrained than constrained grasping. This result is consistent with our hypothesis that S1 provides inputs to M1 for scaling digit forces. We confirmed these hypothetical roles of S1 and M1 through virtual lesions. As anticipated, stimulation of S1 disrupted digit force scaling only in unconstrained grasping. Stimulation of M1 during unconstrained grasping interfered with digit force-to-position modulation and led participants to rely on previously-used digit forces. In contrast, stimulation of M1 during constrained grasping impaired participants’ ability to retrieve forces used in previous manipulations. These results provide comprehensive support for our theoretical framework of dexterous manipulation.

## RESULTS

For both constrained and unconstrained grasp contexts (*con* and *uncon*, respectively), the task consisted of grasping and lifting a sensorized object using the thumb and index fingertip. The task’s goal was to minimize object roll during lift. Participants achieved this goal by exerting a compensatory torque (T_com_) on the object prior to object lift to counteract the object’s external torque (T_ext_) caused by its asymmetrical mass distribution (Fig. 1A; eq. 1, Materials and Methods). As expected from our previous work, we found a significant negative correlation between T_com_ and peak object roll (see Materials and Methods). Therefore, for brevity we focus on T_com_ as measure of anticipatory grasp control and manipulation performance.

**Fig. 1.**
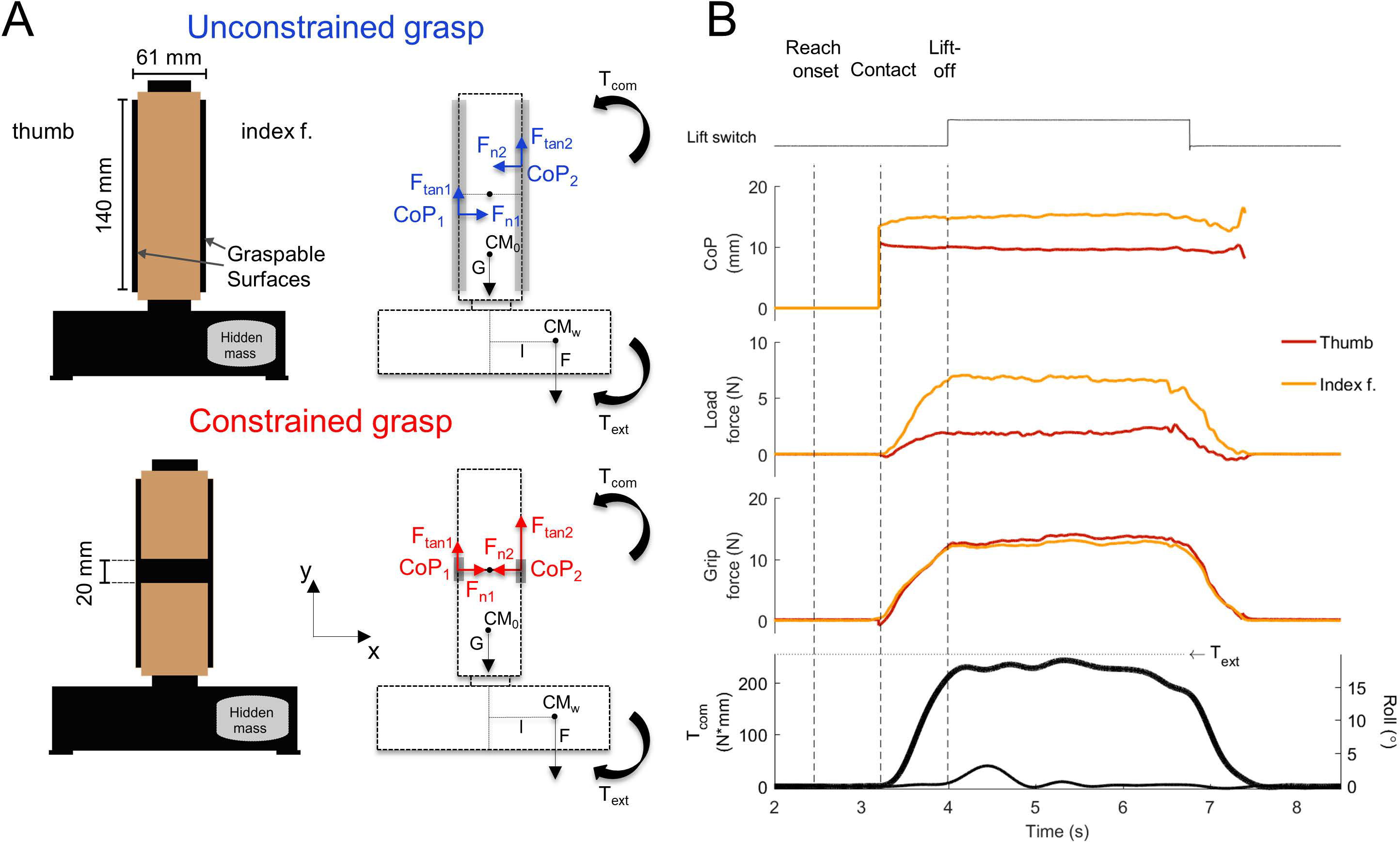
Grip device, experimental conditions, and experimental variables. (*A*) Schematic and free-body diagrams of the custom-built grip device for the unconstrained and constrained grasp conditions. (*B*) Experimental variables are shown for one representative trial of the manipulation task performed using an unconstrained grasp. From top to bottom, traces are thumb and index finger center of pressure (CoP), load and grip forces, compensatory and external torque (T_com_, thick line, and T_ext_, dotted horizontal line, respectively), and object roll (thin line). The sign of T_com_ has been inverted for graphical purposes. At object contact, the index finger is placed higher than the thumb and exerts larger load force. Nearly identical grip force is exerted by each digit. This subject generates a T_com_ that approaches T_ext_ at object lift onset, thus minimizing object roll (thin line; peak value < 5°). Vertical dashed lines from *left* to *right* denote reach onset, contact, and object lift off.

The *con* and *uncon* grasp contexts differed only in terms of whether object contact locations were visually cued or could be chosen by the subject. As expected, we found that subjects in *con* and *uncon* groups learned within the first three trials to generate T_com_. However, digit load forces were significantly correlated with trial-to-trial changes in digit position only for *uncon* grasping (*SI Appendix*, S3). To determine the extent to which the role of M1 and S1 is grasp-context dependent, we conducted electroencephalography (EEG) and transcranial magnetic stimulation (TMS) experiments (Fig. 2A,B).

**Fig. 2.**
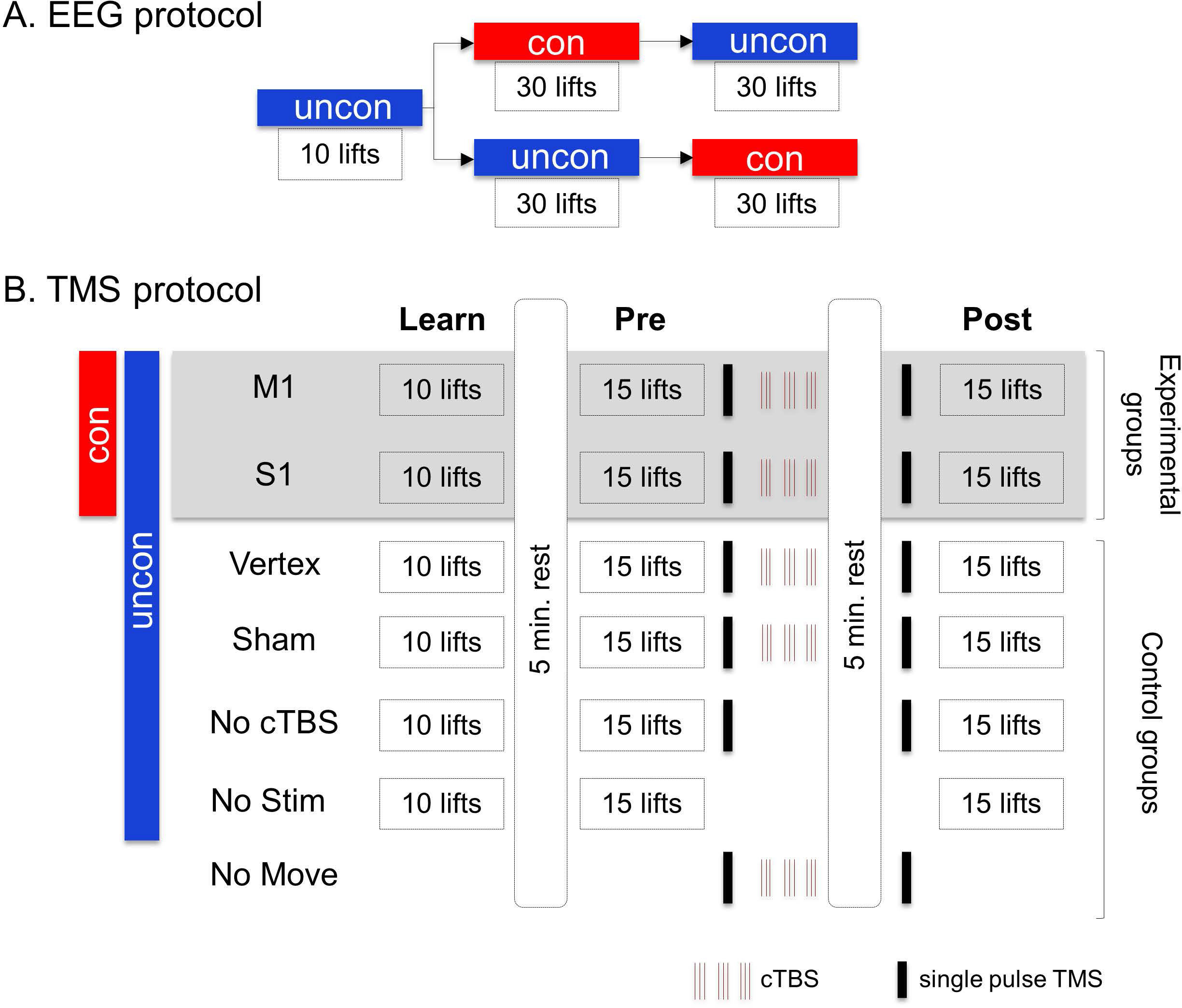
Experimental protocols. (*A*) For the electroencephalography (EEG) experiment, all subjects performed a block of 10 trials of the unconstrained grasp condition (*uncon*), after which they were split in two groups. One of these groups performed 30 trials of the constrained grasp condition (*con*) followed by 30 trials of the unconstrained grasp conditions, whereas the second group performed these grasp conditions in the opposite order. (*B*) In the transcranial magnetic stimulation (TMS) experiments, we delivered single pulse TMS and/or continuous theta burst stimulation (cTBS). The experimental groups (*n* = 4) performed our manipulation task in both *con* or *uncon* grasp conditions and received cTBS over M1 or S1. All control groups (*n* = 5), with the exception of “No Move”, were tested in the *uncon* grasp condition. All groups, with the exception of “No Stim”, received single pulse TMS before and after cTBS. All groups, with the exception of “No Move”, performed 10 repetitions of the manipulation task during the Learn block and 15 repetitions each during the Pre- and Post-cTBS blocks.

### Sensorimotor cortical activity is sensitive to grasp context

The EEG experimental design aimed at removing significant differences in digit position, forces, and torques at object lift onset across the two grasp contexts (see Materials and Methods). This was confirmed by statistical analyses revealing that digit force and position distributions at object lift-off were statistically indistinguishable in *con* versus *uncon* grasping (all *P* > 0.62).

We recorded EEG and quantified source current density power over precentral and postcentral regions (*SI Appendix*, S2). Source power was computed from contact to object lift onset for all trials from both grasp contexts. Importantly, EEG *s*ource power was significantly larger during *uncon* than *con* grasping over both M1 and S1 2 2 (main effect of *Group: F*_1,21_ = 379.6 with 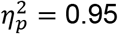 and 22.69 with 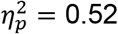, respectively; both *P* < 0.0001; Fig. 3A,B). Furthermore, statistical analysis used to analyze Granger causal directed connectivity between M1 and S1 revealed a significant interaction between the factors of grasp type (*con* versus *uncon*) and connectivity direction (M1 to S1 versus S1 to M1) (F_1,21_ = 3.87, *P* < 0.05, 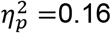). As predicted by our framework, the effective drive was higher going from S1 to M1 during *uncon* compared to *con* grasping (t_21_ = 2.53, *p* < 0.01, Cohen’s d_z_ = 0.54), whereas there was no significant difference in Granger causality from M1 to S1 for *con* and *uncon* grasping (*P* > 0.05) (Fig. 3C). It is worth noting that while changes in power can correlate with changes in connectivity, there is no strict dependency between the two. Therefore, these results indicate the main cortical mechanism differentiating grasp contexts is not an increased site activation, as both M1 and S1 increased in power from *con* to *uncon* (Fig. 3B). Importantly, the connectivity results indicate the main mechanism is an increased information transfer from S1 to M1 in *uncon* grasping.

**Fig. 3.**
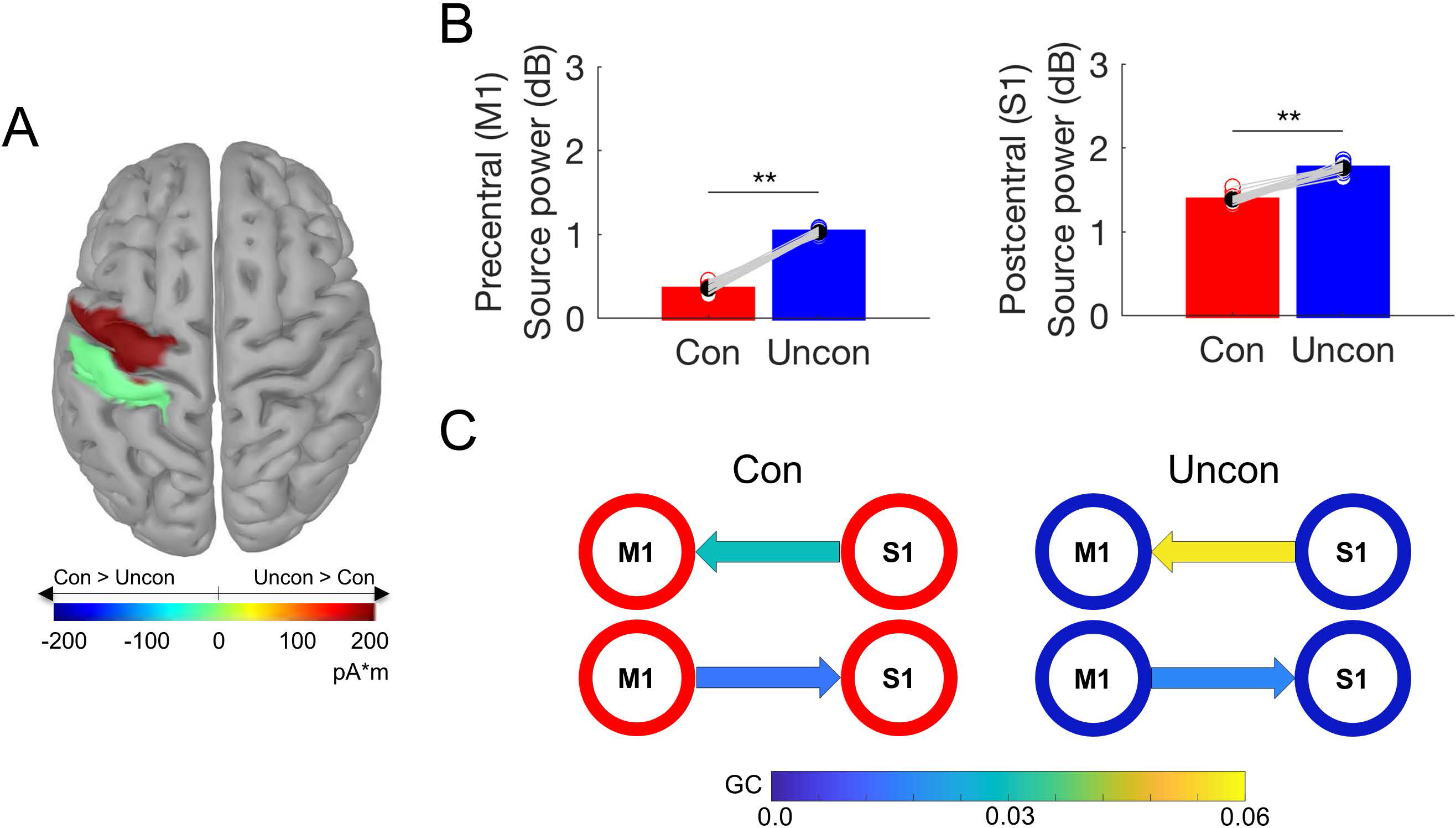
EEG source current density power and effective connectivity. (*A*) Color-coded brain areas denote statistically significant source power differences (pA*m) between constrained and unconstrained grasping over left precentral and postcentral regions. (*B*) Mean normalized source power (decibel) for each grasp context. Overlaid gray lines show condition differences for individual subjects. ** denotes *P* < 0.0125. (*C*) Nodes showing the Granger causal values between M1 and S1 estimated from source reconstructed EEG data. The figure shows nodes for connectivity between M1 and S1 (pre- and post-central gyrus). The color of the arrow between nodes represents the strength of drive between areas. All connectivity values were significantly different than zero (*P* < 0.05). Data in *(A)-(C*) are from all subjects (*n* = 22) and were computed from object contact to lift.

These findings support our hypothesis that the roles and interactions of M1 and S1 are dependent on grasp context, while raising the question of their functional roles. To address the functional roles of M1 and S1 in *uncon* versus *con* grasping, we performed TMS to induce virtual lesions to M1 and S1 using continuous theta burst stimulation (cTBS) (Fig. 2B).

### A virtual lesion of M1 and S1 impairs execution of learned manipulation in a grasp context-specific fashion

On the first trial of the *Learn* block, subjects were unaware of the object’s mass distribution, as the object is visually symmetrical, and therefore exerted negligible T_com_ (Fig. 4A; Learn 1, Fig. 4B). Consistent with previous work (25), all subjects quickly learned to compensate for the object’s mass distribution and generated the necessary T_com_ over the remaining trials of the *Learn* block (main effect of *Block*: *F*_1,45_ = 522.14, *P* < 0.0001, 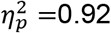 Fig. 4B) similarly across experimental and control groups (no significant *Group* × *Block* interaction: *F*_4,45_ = 2.26 or main effect of *Group: F*_4,45_ = 1.71, both *P* > 0.08). Following learning, T_com_ was stable for all subjects during the remaining *Learn* and *Pre* block trials across experimental and control groups (no significant *Group* × *Block* interaction: *F*_7.85,88.32_ = 0.8, no main effect of *Block*: *F*_1.96, 88.32_ = 2.69, or no main effect of *Group*: *F*_4,45_ = 0.69; all *P*-values > 0.08; Fig. 4B; *SI Appendix*, S5).

**Fig. 4.**
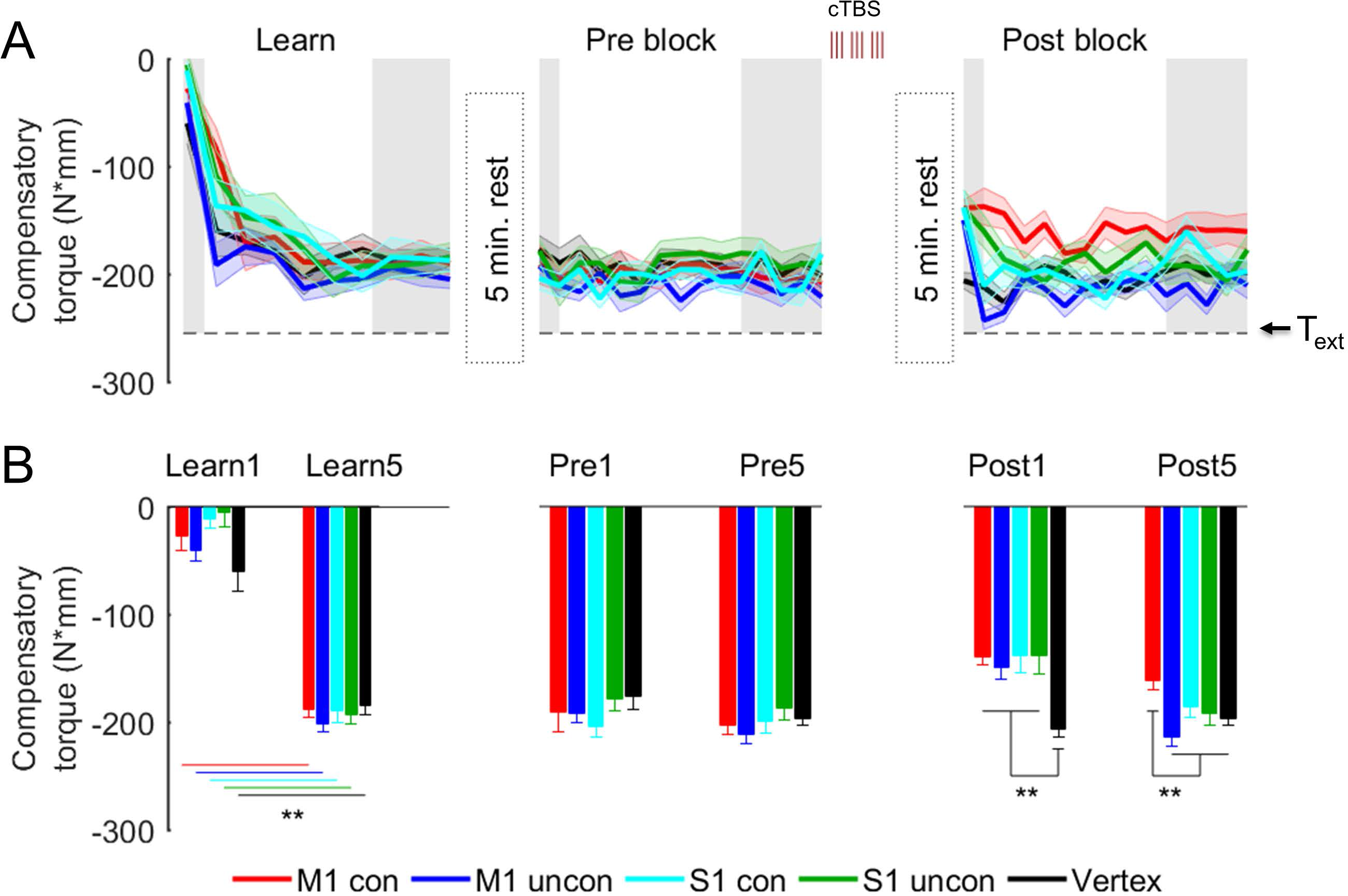
Compensatory torque: Experimental groups and control condition (Vertex). (*A*) Compensatory torque (T_com_) during the Learn, Pre, and Post blocks in the M1 *con*, M1 *uncon*, S1 *con*, S1 *uncon*, and *Vertex* groups. The horizontal dashed line denotes the external torque induced by the added mass at the bottom of the object (T_ext_) that should be compensated for by T_com_. Shaded data denote trials used for plotting in *B* and analysis. (*B*) T_com_ on the first trial and the average of the last 5 trials for each block. ** denotes *P* < 0.0125. Data are averages (± SE) of all subjects.

We delivered cTBS to the M1 *con*, M1 *uncon*, S1 *con*, S1 *uncon*, and *Vertex* groups immediately following the *Pre* block, but prior to the beginning of the *Post* block (Fig. 4). We verified the effectiveness of our cTBS protocol using single-pulse TMS to measure changes in corticospinal excitability in all groups (Fig. 2B;*SI Appendix*, S4 and Fig. S1). We selected the vertex as a neutral control site to assess the specificity of cTBS-induced effects on the control of our manipulation task following stimulation of M1 and S1.

Following cTBS over M1 and S1, but not vertex, subjects were unable to exert the previously-learned T_com_ (significant *Group × Block* interaction: *F*_7.6, 85.53_ = 4.36, *P* < 0.0001, 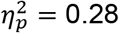 main effect of *Block*: *F*_1.9, 85.53_ = 33.55, *P* < 0.0001, 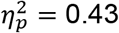) (Fig. 4B). Specifically, on the first trial following cTBS (*Post1*), subjects in all experimental groups exerted significantly smaller T_com_ than those in the *Vertex* group (Dunnett’s t: all *P-*values ≤ 0.004, 1.62 < Cohen’s d < 2.82), although T_com_ reduction was not significantly different across experimental groups (Bonferroni’s t: all *P*-values > 0.99). However, the persistence of the cTBS effect on T_com_ during the *Post* block was dependent on whether subjects performed the manipulation task in the *con* or *uncon* condition and the cortical area targeted by cTBS. For the M1 *uncon*, S1 *uncon*, and S1 *con* groups, T_com_ impairment was short lived, returning to the same magnitude as T_com_ exerted by the *Vertex* group at the end of the *Post* block (Dunnett’s t: all *P*-values > 0.43; Fig. 4B). In contrast, the drop in T_com_ for the M1 *con* group persisted until the end of the *Post* block, as revealed by significantly smaller T_com_ relative to the *Vertex* group (Dunnett’s t: *P* = 0.016, Cohen’s d = 1.12). Similar to the *Vertex* group, no difference in task performance was found in all other control groups when comparing *Pre* and *Post* blocks (*SI Appendix*, S5 and Fig. S2).

Because T_com_ results from the coordination of digit position (d_y_), grip (F_GF_) and load (d_LF_) forces (equation 1, Materials and Methods), we analyzed the extent to which virtual lesions affected each of these variables for each experimental group using the same subsets of trials as those used for the above T_com_ analysis. The similar reduction in T_com_ observed on the first trial following cTBS could have been interpreted as a non-specific effect on T_com_ variables regardless of grasp type and cortical area targeted by TMS. However, the different time courses after cTBS (Fig. 4) indicate that digit position and forces were highly sensitive to the grasp context and cortical area being stimulated.

### Constrained grasping: Virtual lesions of M1 and S1

The M1 and S1 *con* groups learned T_com_ within the first few trials of the *Learn* block by exerting a greater load force on the index finger than thumb (negative d_LF_, equation 1, Methods; this behavior has also been described in previous studies (25, 26, 32)). Furthemore, participants consistently exerted the learned T_com_ throughout the blocks of trials preceding cTBS.

### Disruption of M1 impairs retrieval of learned grip and load forces

Following cTBS, M1 *con* participants were unable to retrieve and use the same digit forces used in previous trials, such retrieval being a key feature of *con* grasping (19, 33). This effect of cTBS started ~200 ms after contact, leading to significantly smaller digit forces by the time the object was lifted (Fig. 5A, M1 *con* column) (main effect of *Block*: *F*_2,18_ = 14.30 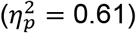, d_LF_ and 13.76 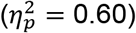, F_GF_; both *P* < 0.0001). d_LF_ and F_GF_ on the first *Post* block trial were significantly smaller than on the late *Pre* block trials (post1 vs. pre5: t_9_ = −7.081 (Cohen’s d_z_ = 2.24) and 3.936 (Cohen’s d_z_ = 1.24), respectively; both *P* < 0.003; Fig. 5A). Importantly, and in contrast to digit forces, d_y_ was unaffected by cTBS (no main effect of *Block*, *F*_2,18_ = 0.045, *P* = 0.956; 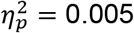; Fig. 5A,B). These results indicate that the effect of cTBS on T_com_ in *con* grasping (Fig. 4) was due to selective disruption of the retrieval of learned digit forces, while sparing the control of visually-cued digit position. Note that the effects of cTBS on digit forces persisted throughout all *Post* block trials (Fig. 5B; *SI Appendix*, S6 and Fig. S3).

**Fig. 5.**
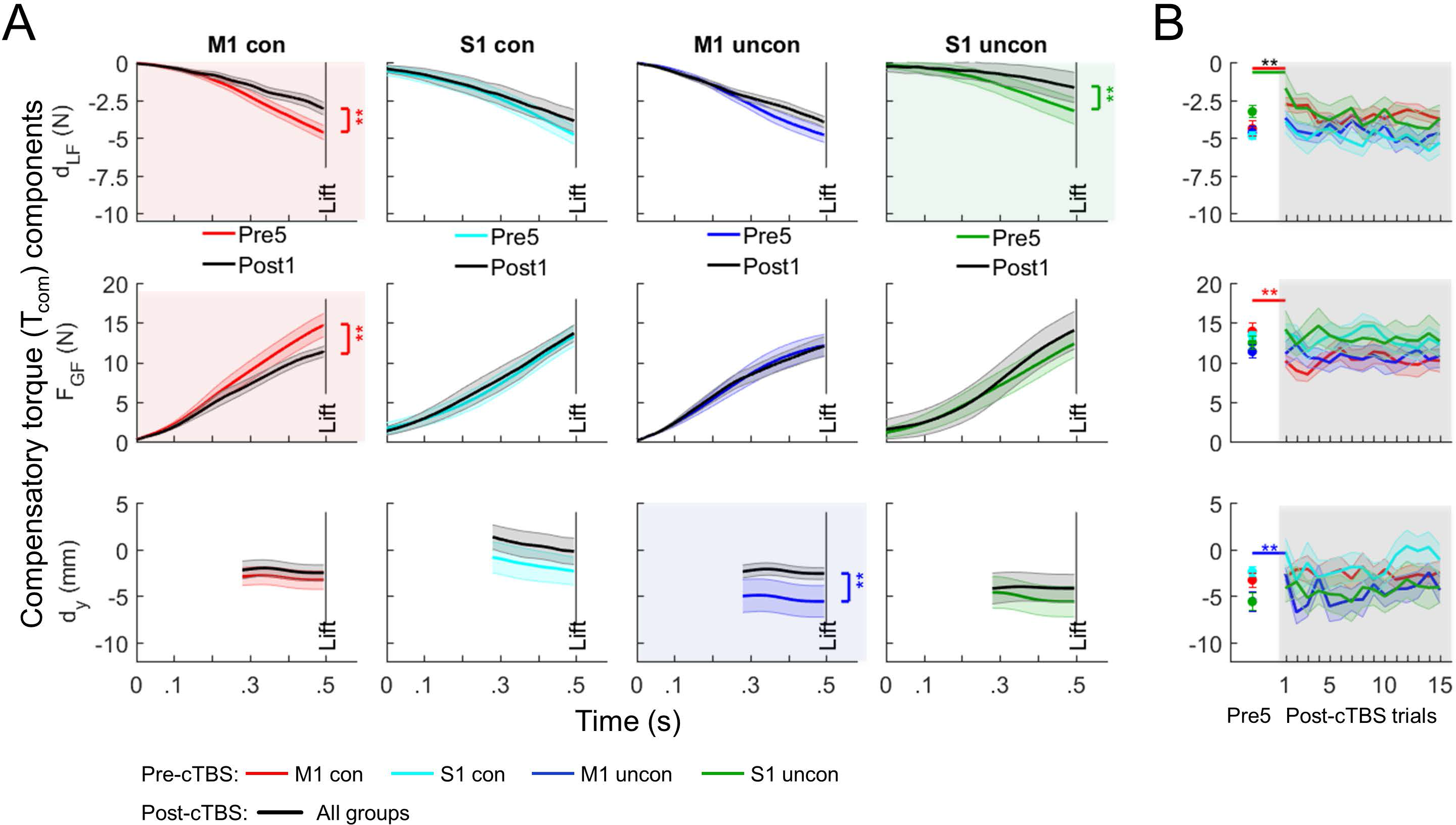
Effect of cTBS on digit load force, grip force, and position across all experimental conditions. (*A*) From top to bottom, traces denote time course of the difference between thumb and index finger load force (d_LF_), grip force averaged across thumb and index finger (F_GF_), and the vertical distance between thumb and index finger center of pressure (d_y_) from contact (“0”) to object lift onset. Data are averages of the last 5 trials prior to cTBS (Pre5) and first trial following cTBS (Post1). d_Y_ data are plotted from the time at which they can be accurately estimated using force and torque sensors (25). Data from each experimental group are shown across columns. Shaded plots denote T_com_ variables that were significantly affected by cTBS. (*B*) Data from Pre5 and each post-cTBS trial are shown for each T_com_ variable and experimental group. ** denotes P < 0.0125. Data are averages (± SE) of all subjects.

### Disruption of S1 does not impair the modulation of digit forces or position

Unlike the M1 *con* group, none of the T_com_ components was affected by cTBS in the S1 *con* group (no main effect of *Block*; d_LF_: *F*_1.3, 11.74_ = 2.6, F_GF_: *F*_2,18_ = 2.2, and d_Y_: *F*_1.8, 16.5_ = 3.5 respectively; all *P*-values > 0.05; Fig. 5A, S1 *con* column). These results indicate that the effect of cTBS of S1 on *con* grasping was not due to a significant disruption of the retrieval of learned digit forces or the control of visually-cued digit position. Therefore, the reduction in T_com_ on the first post-cTBS trial (Fig. 4) was caused by a small, non-significant effect on d_Y_ and its multiplicative effect on F_GF_ contribution to T_com_ (equation 1, Materials and Methods).

### Unconstrained grasping: Virtual lesions of M1 and S1

Consistent with our previous behavioral work (25, 26, 32), M1 and S1 *uncon* groups learned to exert T_com_ to counter the clockwise T_ext_ by exerting greater load force with the the index finger and placing it higher than the thumb (negative d_LF_ and d_Y_ respectively; equation 1, Methods). Unlike *con* grasping, d_LF_ and d_y_ significantly covaried across the *Learn* and *Pre* block of trials (*SI Appendix*, S3).

### Disruption of M1 impairs control of learned digit position and modulation of load force

cTBS to M1 impaired subjects’ ability to use similar digit positions learned in previous *uncon* trials (main effect of *Block*: *F*_2,18_ = 10.29, *P* = 0.001, 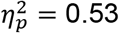). Specifically, on the first post-cTBS trial the vertical distance between thumb and index finger center of pressure (d_Y_) significantly decreased relative to the late *Pre* block trials (post1 vs pre5: t_9_ = −4.384, *P* = 0.002, Cohen’s d_z_ = 1.39) (Fig. 5A, M1 *uncon* column). Note that the large change in d_Y_ caused by cTBS was not accompanied by a significant modulation of d_LF_ or F_GF_ (no main effect of *Block*: *F*_2,18_ = 5.27 and 0.106, respectively; both *P* > 0.05; both 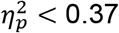). This is an important observation, given that d_LF_ modulation to d_Y_ is a key feature of *uncon* grasping which was found during *Learn* and *Pre Block* trials. Thus, the effects of cTBS during *uncon* grasping were opposite to those found for *con* grasping: virtual lesion to M1 impaired the control digit placement, but not digit forces. These results indicate that the lack of modulation of d_LF_ to the cTBS-induced change in d_Y_ caused the drop in T_com_ in the early trials of the *uncon* grasping condition (Fig. 4A). However, on subsequent trials a strong modulation of d_y_ and concurrent modulation of d_LF_ enabled T_com_ to return to pre-cTBS levels (Fig. 5B; *SI Appendix*, S6 and Fig. S3).

### Disruption of S1 impairs the modulation of load force distribution

cTBS over S1 affected only digit load force distribution, d_LF_ being significantly reduced relative to *Pre* block trials (main effect of *Block*: *F*_2,18_ = 16.50, *P* < 0.0001, 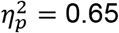; post1 vs pre5: t_9_ = –4.187, *P* = 0.002, Cohen’s d_z_ = 1.32; Fig. 5A,S1 *uncon* column). In contrast, F_GF_ and d_y_ were statistically indistinguishable from trials preceding cTBS (no main effect of *Block*: *F*_2,18_ = 0.867 and 2.34, respectively; both *P* > 0.13; both 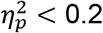; Fig. 5*A*). Therefore, the reduction in d_LF_ was the primary cause of T_com_ reduction on the first post-cTBS trial (Fig. 4A). Subsequent modulation of d_LF_ contributed the most to the recovery of T_com_ to pre-cTBS levels (Fig. 5B; *SI Appendix*, S6 and Fig. S3).

## DISCUSSION

Dexterous manipulation requires not only precise force application over time, but also adjusting forces to where they are applied on an object. In most every day hand-object interactions, digit contact points are not cued or constrained by an object’s geometrical features. Nevertheless, humans are exquisitely skilled at repeatedly manipulating an object despite the fact that grasping may occur at different contact points on the object. Such variability in digit placement persists even after the object’s dynamic properties have been learned (25). Digit position variability is compensated for by modulating digit forces in an anticipatory fashion, i.e., before object lift-off (25, 26). This digit force-to-position modulation is critically important for ensuring that manipulation goals are accurately accomplished. This evidence points to the involvement of a feedback-based control of digit forces. In this case, sensed digit placement would be used to modulate digit forces accordingly on a trial-to-trial basis.

From a neural control perspective, we had proposed that constraining or allowing choice of contact points challenges the nervous system in different ways (25–27, 30, 34). Constraining digit position enables the memory retrieval of grasp forces used in previous manipulations. In contrast, the larger digit position variability when allowing choice of contacts limits the extent to which subjects can use memory-based control of forces. The present study tested whether digit force-to-position modulation is mediated by differential activation and communication between sensorimotor cortices.

Our combined results across EEG and TMS studies revealed a differential involvement of M1 and S1 in unconstrained relative to constrained grasping. While we demonstrated that the neural activity of both M1 and S1 is greater in unconstrained grasping, only the effective connectivity from S1 to M1 was affected by grasp context (Fig. 3C). A series of TMS experiments further elucidated the functional roles of M1 and S1. When digit position was constrained to be repeatable and predictable across trials, we confirmed previous work pointing to the role of M1 in mediating a memory-based control of manipulative forces (15, 17) and S1’s lack of involvement in digit force retrieval (31) (Fig. 5A, M1 *con* and S1 *con*). More importantly, for unconstrained grasping we demonstrated that integrity of both M1 and S1 is critically important as they have complementary roles in mediating digit force-to-position modulation (Fig. 5A, M1 *uncon* and S1 *uncon*). Together, our results suggest that control of dexterous manipulation relies on a flexible organization of the sensorimotor cortical network depending on whether contact points can be chosen or not.

### Effects of cTBS on grasp control variables are sensitive to grasp context

Previous work combining TMS-induced virtual lesions and a constrained grasping task identified M1 as being implicated in storage and retrieval of sensorimotor memory of grasp forces (15, 17, 35). Our results confirm such a role, while extending these observations in important ways. Specifically, when contact points were predictable, virtual lesions to M1, but not S1, prevented retrieval of learned digit forces (M1 and S1 *con*, Fig. 5A). In contrast, when contact points were not as predictable as in constrained grasping, virtual lesions to M1 elicited two inter-related phenomena: subjects could not implement digit placement similar to that before delivery of cTBS, and digit forces were not modulated as a function of the new digit placement (M1 *uncon*, Fig. 5A).

Furthermore, cTBS to S1 impaired digit force-to-position modulation (M1 *uncon*, Fig. 5A). Thus, in both *uncon* groups, cTBS impaired the critical ability to modulate digit forces to position, but did so by selectively affecting different T_com_ variables.

Analysis of the time course of cTBS effects beyond the first post-cTBS trial provided additional insights. Specifically, when digit force control was predominantly driven by sensorimotor memory, the M1 *con* group’s inability to retrieve digit forces and restore pre-cTBS T_com_ persisted for all post-cTBS trials (Fig. 5B). In contrast, for the S1 *con* group the small (non-significant) effect of cTBS on d_Y_ (Fig. 5A) and T_com_ reduction disappeared after the first post-cTBS trial due to small changes in digit centers of pressure (Fig. 5B; *SI Appendix*, S6 and Fig. 3S). Importantly, the M1 and S1 *uncon* groups were able to restore digit force-to-position modulation and T_com_ within the first five post-cTBS trials.

The different time courses of post-cTBS recovery in each T_com_ variable further indicates differences in the roles of M1 and S1 according to the grasp context. The most striking difference was found in the timeline of post-CTBS effects across M1 and S1 *con* groups, i.e., F_GF_ and d_LF_ were affected for 15 trials, whereas the small effect on d_Y_ lasted 1 trial, respectively (Fig. 5B; *SI Appendix*, S6 and Fig. 3S). These findings confirm M1 – but not S1 – is involved in storing or retrieving memory of digit forces. As cTBS to S1 did not affect digit forces, the quick recovery of d_Y_ through small changes in digit position in the S1 *con* group could have been driven by visual feedback of object roll caused by the sudden T_com_ reduction on the first post-cTBS trial (Fig. 4A,B). These results suggest that the memory-based force control mechanism affected by cTBS cannot benefit from visual feedback of manipulation error to the same extent as digit placement, even when such errors continue to occur across multiple trials.

With regard to the *uncon* groups, cTBS to M1 and S1 again affected different T_com_ variables, i.e., d_Y_ and d_LF_, respectively (Fig. 5A). Importantly, both groups were able to restore pre-cTBS T_com_ by re-establishing digit force-to-position modulation, but did so in different ways. Specifically, the M1 *uncon* group modulated both d_Y_ and d_LF_, whereas the S1 *uncon* group modulated only d_LF_ (Fig. 5B; *SI Appendix*, S6 and Fig. 3S). These differences in short- and long-term effects of cTBS, as well as the T_com_ variables affected by the virtual lesion, underscore the complementary, yet different roles of M1 and S1 in unconstrained grasping. Specifically, integrity of both M1 and S1 is needed to modulate digit force to position, even though a virtual lesion to either area can be compensated by the other. Similar to the recovery of pre-cTBS levels of T_com_ proposed for the *con* groups, we speculate that this recovery in the *uncon* groups was also mediated by visual feedback of object roll, as well as digit placement which could be changed in a trial-to-trial basis.

The above results contribute to extant evidence indicating M1 is largely involved with motor memory processes. For example, previous studies have shown that neurostimulation to M1 during motor learning tasks affects recall but not the learning process itself (36, 37). Importantly, in unconstrained grasping cTBS to M1 disrupted digit force modulation to position, while leading participants to rely on memory of forces (Fig. 5A). These findings indicate the role of M1 is not limited to a memory-based control process: it is also directly involved in using trial-by-trial sensory feedback of digit position to scale forces.

Our focus on S1 was motivated by a long history of research on the role of S1 in the context of online feedback control (e.g., (38–40)). While it is clear that S1 provides proprioceptive inputs to M1 (41–44), what remains unclear is whether this process is context dependent. We hypothesized that increased communication of proprioceptive inputs from S1 to M1 would be necessary in unconstrained grasping. Our cTBS results support this prediction by showing an impaired ability to adjust forces online to position. This suggests that S1 cTBS inhibited communication to M1 and supports S1 has a context-dependent role. Recent evidence further supports this proposition by indicating that S1 is also involved in motor learning (45). These authors demonstrated that optogenetically silencing S1 during motor adaptation inhibited learning. The implication of this finding and our S1 cTBS results is that somatosensory cortex plays a critical role in providing feedback information for both online control and learning.

### M1-S1 communication specificity across grasp contexts

The cTBS results highlight the individual roles of M1 and S1 as a function of grasp context. Nonetheless, they clearly show that their functional roles are not independent. Our framework predicted S1 would increases communication of proprioceptive inputs to M1 in unconstrained grasping. We used EEG connectivity analysis to test this hypothesis. Specifically, we anticipated a two-way interaction between grasp context and the extent to which S1 drove activity in M1, and vice versa. Our analysis confirmed this prediction that connectivity from S1 to M1 was increased in unconstrained grasping (Fig. 3B,C). Accordingly, greater connectivity from S1 to M1 in unconstrained grasping and the behavioral effect of S1 disruption are consistent with the proposition that S1 would provide M1 with feedback of digit position. This raises an auxiliary but important question of which brain area may set the context-dependency. Previous work on M1-S1 interactions in rats concluded that M1 may cue S1 to gate thalamic inputs in response to whisker stimulation (46). We found that M1 to S1 connectivity was similar across grasp contexts, suggesting that the control of sensory gating may also be driven by S1.

Beyond connectivity, an imperative question is whether local activity in M1 and S1 was also context dependent. When examining power, we focused our analysis on 9-12 Hz (alpha). We chose this frequency band because substantial support has been provided for its association with cortical inhibition within sensory cortices (47). Unlike the connectivity results, we found that power increased in both M1 and S1 in unconstrained grasping. We suggest that the increase in power likely represents an increase in local inhibition across both areas to gate inputs from external sources (48, 49). This overall increase in power, combined with the connectivity result, indicate that both local inhibition and inter-areal communication may instatiate distinct processes during feedback-driven sensorimotor control.

### Primacy of contact event for somatosensory feedback

We should note that S1’s role in processing sensory inputs associated with digit position is not obligatory. Conceptually, force planning could arise prior to contact, when the digits are visible. While such vision-based force planning is still possible, contact detection through visual, proprioceptive, and tactile feedback has been shown to be a critical event for signaling the transition from the end of hand transport and onset of force application (for review see 11). We propose that feedback during object contact is also instrumental for estimating the relative position of the digits. Importantly, the S1 *uncon* results indicate that visual feedback of the hand trajectory and contact points – available throughout the task – could not compensate for the effect of the virtual lesion on digit force-to-position modulation. The notion that object contact is the most relevant event for feedback processing in unconstrained grasping is supported by a recent study showing significant grasp-context differences in corticospinal excitability at contact, but not during the reach (30). In summary, our findings support the imperative role of somatosensory feedback of digit position for digit force modulation at object contact, but not during reaching.

### Grasp cortical network for constrained and unconstrained grasping

It is well known that the cortical network underlying grasp control, which has been defined primarily based on research on constrained grasping, comprises several areas. For example, the anterior region of intraparietal sulcus (aIPS) and premotor ventral (PMv) contribute to accurate hand shaping prior to object contact (8), whereas aIPS, premotor dorsal area (PMd), and M1 are involved in the storage and/or retrieval of force scaling appropriate for the hand shape used in the grasp (50). Studies in patients with posterior parietal cortex lesions revealed that they experience difficulties in shaping the fingers according to intrinsic object features (51) and impaired predictive scaling of grip forces during self-induced modulation of load forces (52). In neurologically-intact individuals, TMS-induced disruption of PMv and bilateral aIPS areas prior to object lift-off leads to increased variability in hand shaping (14, 53–55), whereas TMS disruption of contralateral aIPS, PMd and M1 results in inaccurate grip force and load force scaling (14, 15). S1 appears to be involved in sensing of predictable (31) and unpredictable (17) contact events occurring at the fingertips during constrained grasping. In sum, when contact points are constrained, dexterous manipulation relies on interactions among posterior parietal areas, such as aIPS and S1, and frontal areas such as PMv, PMd, and M1 (56, 57).

Besides the above-cited study on corticospinal excitability (30), to the best of our knowledge only one study examined the extent to which control of constrained and unconstrained grasping is mediated by different brain areas (29). The main findings of this fMRI study were that cerebellum, BA 44, and PMv were differentially activated across the two grasp contexts. However, in contrast to our EEG and cTBS results, no differences were found in M1 and S1 activity. A direct comparison between these studies is not possible due to several methodological differences, i.e., in the fMRI study T_com_ variables could not be measured in the scanner and subjects were instructed to deliberately vary digit position across trials in the unconstrained condition. Nevertheless, in the present study our cTBS results identified a causal and grasp context-dependent role of M1 and S1, which is consistent with the EEG results of greater S1-M1 effective connectivity. Future studies are needed to extend these observations to other brain areas of the grasp network.

### Revised conceptual framework of neural control of manipulation

The present findings significantly extend our understanding of neural mechanisms underlying dexterous manipulation. Specifically, they reveal that the neural and control mechanisms are differentiated between constrained and unconstrained grasping, the latter mimicking more natural conditions. These findings necessitate a revision to current frameworks explaining dexterous manipulation. We used the current and previous results to provide the foundation for a revised theoretical framework (Fig. 6).

**Fig. 6.**
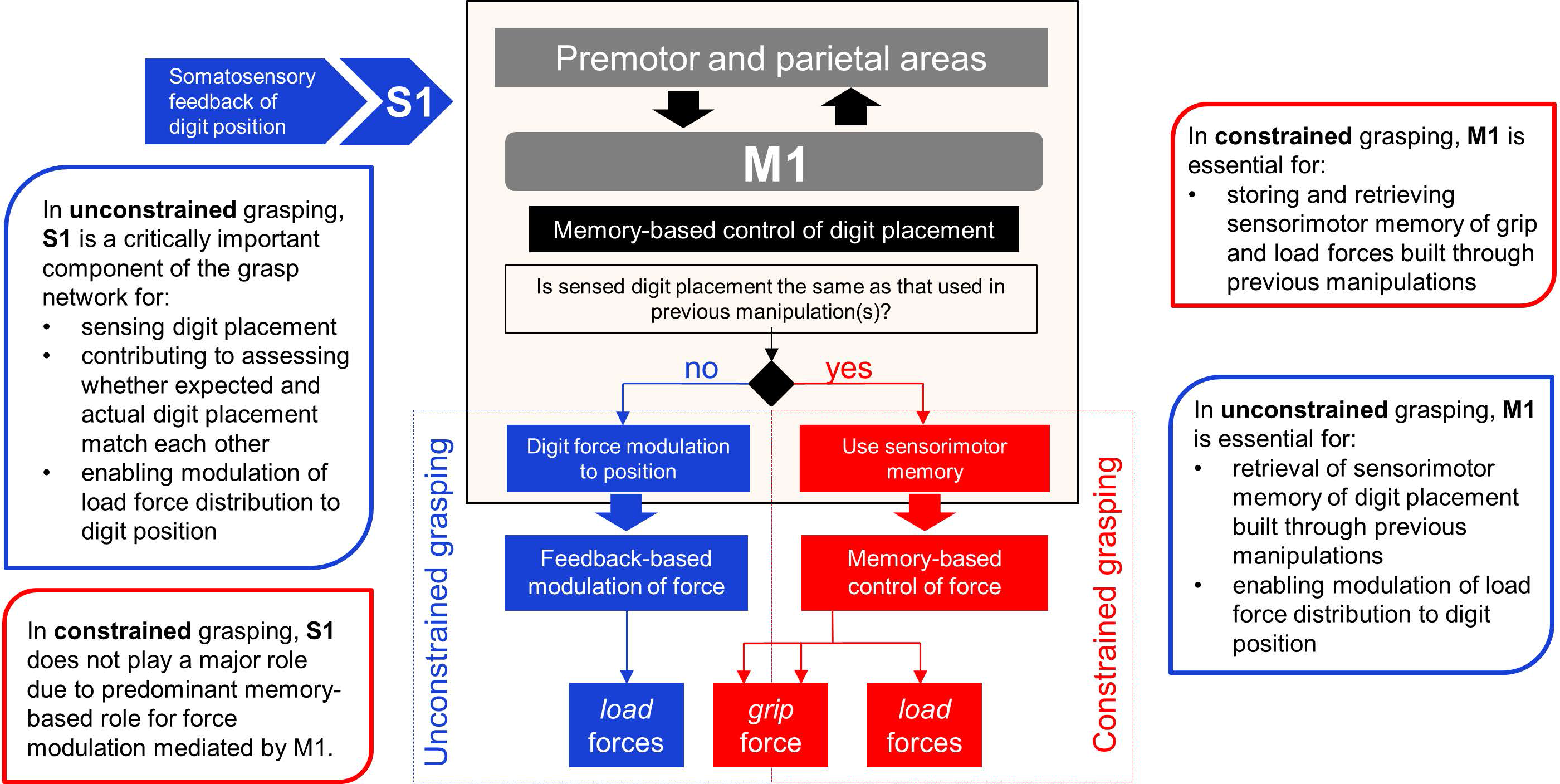
Cortical sensorimotor mechanisms for neural control of dexterous manipulation. The diagram shows a revised model describing the roles of M1 and S1 for the control of dexterous manipulation for constrained and unconstrained grasping contexts.

It has been established that interactions between M1, sensory, as well as premotor and parietal cortical areas, lead to hand shaping (58, 59), which culminates with positioning the digits at remembered locations used in previous manipulations. Somatosensory and visual inputs contribute to guiding the hand towards the planned contact points on the object (Fig. 6).

Our theory posits that following object contact, subjects use feedback of digit position to determine the similarity of contact points with those used in previous manipulations. We propose that sensing digit position serves a dual role. The first and obvious role is detecting the digit contact points. However, the utility of this is driven by whether these contact points were constrained or unconstrained, necessitating a second role. For constrained grasping, this role is minimally important beyond ensuring force control that satisfies mechanical requirements, i.e., normal-to-load force modulation (33, 60). In contrast, during unconstrained grasping we propose that manipulation is predominantly driven by a mechanism that compares predicted and actual sensory feedback of digit position (Fig. 6). We anticipate that individuals use this mechanism to determine the extent to which force control is driven by memory and online feedback. The relative contribution of each mechanisms depends on the extent to which predicted and actual contact points match. Daily grasping activities, e.g., unconstrained grasping, elicit greater reliance on feedback because of the inherent variability of contact points.

Our work underscores the context-dependent roles of M1 and S1 for the control of dexterous object manipulation by revealing the dual role of M1 and its synergistic interactions with S1 for the trial-to-trial modulation of digit force to position. Importantly, adding one degree of freedom (choice of digit position) to the control space of manipulation elicited *different* roles of M1 and S1 to perform the *same* task. Specifically, as the number of solutions (force-position relations) increases, the communication between M1 and S1 becomes more critical. Future work should examine the contribution of higher-order frontal and parietal cortical areas to building high-level task representations required to drive the coordination of digit position and forces to attain a given task goal, i.e., preventing the object from slipping or tilting during manipulation.

## MATERIALS AND METHODS

### Subjects

One hundred naïve right-handed volunteers (23 ±4.12 years [mean ± SD]; 44 females) with normal or corrected-to-normal vision and no history of musculoskeletal disorders or neurological disease participated in this study. Subjects were screened for potential risks of adverse reactions to transcranial magnetic stimulation (TMS) using the Transcranial Magnetic Stimulation Adult Safety Screen (61) and gave their written informed consent according to the declaration of Helsinki. All protocols were approved by the institutional review boards at Arizona State University and the University of Houston.

### Grip device

A custom-designed inverted T-shaped object instrumented with two six-dimensional force and torque transducers (Nano 25; ATI Industrial Automation, Garner, NC) (Fig. 1*A*) was used to record forces and torques exerted by the index fingertip and thumb. Graspable surfaces (sandpaper, grit #320) consisted of two long parallel PVC plates (140 × 22 mm), each mounted vertically on one transducer (Fig. 1*A*). The grip device measured grip and load force (normal and tangential/vertical to the graspable surface), and each digit’s center of pressure. The transducers’ location relative to the graspable surfaces was blocked from the subject’s view to prevent visual cues from biasing the choice of digit placement. A 400-g mass was placed in the right (relative to the subject) compartment at the base of the grip device and was hidden from view to prevent subjects from anticipating the object’s mass distribution. The added mass created an external torque in the frontal *x-y* plane of 255 N•mm (T_ext_, Fig. 1*A*). The object’s total mass was 790 g. Each end of the object’s base was placed on a lift switch. The release of either switch by upward movement of the object from the table signaled object lift onset. We used a wireless inertial measurement unit (IMU, Emerald, APDM, Portland, OR) fastened to the top of the object to measure object tilt during the lifting phase.

### Experimental Protocol

Subjects sat comfortably in a custom TMS chair (Rogue Research Inc., Canada), with right hand pronated and resting on a hand switch. Subjects were asked to reach and grasp the grip device placed on a table 15 cm in front of them using the thumb and index fingertip at a self-selected speed, lift the object vertically to a height of 5–10 cm above the table, hold the object for 2–3 seconds, replace the object on the table, and return their hand to the hand switch until the next trial. During each trial, subjects were asked to lift the object as straight as possible, i.e., to prevent the object from rotating on the frontal plane due to the right-sided asymmetrical mass distribution (Fig. 1*A*). Successful performance required subjects to exert a compensatory torque (T_com_) of the same magnitude but in the opposite direction of T_ext_ in an anticipatory fashion, i.e., at object lift onset (25). The rationale for the task design is described in S1 (*SI Appendix*).

Subjects were asked to perform our manipulation task by either allowing them to choose grasp contact locations (unconstrained grasping, *uncon*) or constraining contact locations by visually cueing grasp points on the object (constrained grasping, *con*) (top and bottom objects, respectively, in Fig. 1*A*). The functional roles of M1 and S1 underlying control of *con* and *uncon* grasping were investigated in two separate experiments using electroencephalography (EEG) or transcranial magnetic stimulation (TMS).

For the *uncon* grasping condition in both EEG and TMS experiments, subjects were instructed that they could grasp anywhere along the vertical plates to perform the task. For the *con* grasping condition in the EEG experiment, the contact point of each digit was visually cued using two LEDs (vertical distance: 14 mm) on each side of the object within which the fingertip had to be placed. The EEG study addressed differences in source activation of primary motor and sensory cortices (M1 and S1, respectively) and their effective connectivity during execution of *con* versus *uncon* grasping. We expected these differences to reflect a greater involvement of feedback-vs. memory-based control of forces in *uncon* than in *con* grasping (27). We asked subjects to perform a learning block of 10 *uncon* trials (Fig. 2*A*). As the largest performance improvements (i.e., minimization of peak object roll) occurs within the first three trials (25, 26), we used the mean of subjects’ preferred digit placement at object lift onset in trials 6-10 of this learning block to set the location of the LED boundary locations for the *con* context. This procedure ensured that average digit position and force distributions would be statistically indistinguishable across *con* and *uncon* trials, while leaving intact the trial-to-trial force position modulation which only occurs in the *uncon* grasping condition. Thus, EEG activation differences that might occur when comparing *con* versus *uncon* grasping conditions would only be attributable to processes associated with digit force-to-position modulation occurring between contact and object lift-onset in the *uncon* grasping condition. After the learning block, subjects performed 60 experimental trials that were used for EEG analysis. Specifically, half of the subjects performed a block of 30 *uncon* trials followed by a block of 30 *con* trials, whereas the other half performed these two blocks of trials in the opposite order (Fig. 2*A*). This design controlled for potential order effects of block presentation. Statistical analysis confirmed lack of significant differences in T_com_ and related variables (see equation 1) within each grasping condition regardless of the order of presentation.

Unlike the EEG study, the *con* grasping condition in the TMS experiment was designed to address the question of whether the control of digit force and position differed following a virtual lesion to M1 (Fig. 2*B*). Specifically, the pre-versus post-TMS comparison was performed within each grasping condition, and therefore did not require to statistically match digit force and position at object lift onset across *con* and *uncon* grasping conditions. To allow comparison with previous studies of *con* grasping, subjects were instructed to grip the object at fixed collinear locations indicated by a horizontal marker placed across the front of the object (vertical length: 20 mm; see Fig. 1*A*). For both *uncon* and *con* grasping conditions, a computer monitor placed behind the object presented two visual cues to the subject to guide each trial. The first ‘ready’ cue signaled the beginning of a trial, and after a random delay (1-3 s) subjects were shown a ‘go’ cue to initiate the reach. To allow subjects to learn the dynamics of the object, they were asked to perform 10 practice trials (“*Learn”* block) (Fig. 2*B*). Following this block, subjects then performed two blocks (*“Pre”* block, *“Post”* block) of 15 trials each. TMS was delivered between the *Pre* and *Post* block (see below). Each block was separated by a rest time of 5 minutes (Fig. 2*B*).

### Electroencephalography: Procedures

EEG was recorded from 22 subjects using a 64-channel Acticap system (BrainVision, Morrisville, NC) at a sampling rate of 1 kHz, with resolution 0.1 μV and bandpass filter of 0.1-100 Hz, and impedances kept < 10kΩ. 3-D electrode locations were recorded using a Captrak camera system (BrainVision, Morrisville, NC). EEGLAB was used to perform EEG pre-processing steps (62). Continuous data were first high-pass filtered (0.5 Hz). Eye movement and blink artifacts were removed using Independent Components Analysis (Extended INFOMAX algorithm)(63). On average, 4.46 (± 3.33) components were removed per participant. General EEG procedures are described in S2 (*SI Appendix*).

### Transcranial magnetic stimulation: Procedures

We delivered single-pulse TMS (spTMS) to primary motor cortex (M1) of 78 subjects using a Rapid^2^ stimulator (Magstim, 70-mm figure-of-eight coil, Whitland, UK). Suprathreshold TMS pulses delivered over contralateral (left) M1 representing the right first dorsal interosseus muscle (FDI) were used to estimate resting motor threshold (rMT) (64). To assess corticospinal excitability (CSE), we delivered spTMS with the intensity set at 120% of rMT over the identified FDI region.

We delivered continuous theta burst stimulation (cTBS) to M1 and S1 at an intensity of 80% of active motor threshold (aMT) (65) to transiently disrupt neural activity. Repetitive pulses were delivered in the form of 3 pulses at 50 Hz repeated every 200 ms for 40 s (600 pulses) (65, 66). As cTBS over M1 has been shown to decrease the size of MEPs, we measured CSE using spTMS to verify the effects of cTBS over M1 (66) and S1 (67).

For M1 cTBS, the TMS coil was positioned over the left cerebral hemisphere representing the right FDI muscle, as identified during rMT estimation. For S1 cTBS, we positioned the TMS coil over the postcentral gyrus posterior to the M1 FDI hotspot (68). To locate the stimulation site, we used high-resolution T1-weighted MRI scan (3T Philips Ingenia scanner) obtained from each subject and used it to reconstruct a three-dimensional brain to display the cortical surface (Brainsight software, Rogue Research Inc., Canada) (65). The mean Montreal Neurological Institute coordinates of the stimulation sites for left S1 were −41.75 ±10.37, −25.27 ±15.21, 57.11 ±5.47 (x, y, z, mean ±SD; *n* = 20). For vertex stimulation (see below), the TMS coil was positioned over Cz, based on the 10-20 international system (69) with the TMS handle oriented posteriorly in alignment with the interhemispheric fissure (70). The coil position for S1 and vertex was confirmed by the delivery of single TMS pulses at 120% of rMT to ensure that there were no MEPs in the FDI muscle. General TMS procedures are described in S2 (*SI Appendix*).

### TMS experiment: Experimental Groups

We delivered cTBS to four groups of subjects (Fig. 2*B*). We stimulated M1 and S1 of subjects performing the *con* and *uncon* grasping condition (*M1 con*, S1 *con*, M1 *uncon*, and S1 *uncon*; *n* = 10 in each group). cTBS was delivered between the *Pre* and *Post* blocks. CSE was assessed using spTMS immediately before and 5 minutes after cTBS (66) (Fig. 2*B*).

### TMS Experiment: Control Groups

We performed five control experiments to assess the specificity of cTBS effects over M1 and S1 and the efficacy of the cTBS protocol: *Vertex* (*n* = 10), *Sham* (*n* = 10), *No cTBS* (*n* = 6), *No Stim* (*n* = 6), and *No Move* (*n* = 6). All control groups, with the exception of the *No Move* group, performed the manipulation task in the *uncon* grasping condition (Fig. 2B). Unless otherwise stated, cTBS occurred between the *Pre* and *Post* blocks, and CSE was assessed over contralateral (left) M1 region immediately after the *Pre* block and before the *Post* block (Fig. 2*B*).

In the *Vertex* group, cTBS was delivered over vertex to assess specificity of cTBS-induced effects observed in the M1 and S1 groups (65, 71).

In the *Sham* group, cTBS was delivered using a second coil placed directly behind the TMS chair’s headrest with current directed away from the scalp while the coil over contralateral (left) M1 remained in place. This group was used to control for any somatosensory effects caused by the auditory cue of cTBS on the control of object manipulation (72).

For the *No cTBS* group, no cTBS stimulation was used. This served to quantify the potential effects of MEP-induced movements on manipulation performance, as these muscle twitches have been shown to affect grasping behavior (35).

Subjects in the *No Move* group received cTBS over contralateral M1, and saw the same visual cues as those presented to all other groups. However, they were asked to remain at rest when seeing the ‘go’ cue rather than performing the motor task. This control group was used to validate the effects of cTBS over M1 on MEP size using the protocol that has been previously reported in the literature (66).

Subjects in the *No Stim* group received neither spTMS nor cTBS (Fig. 2*B*). This group was used to control for any somatosensory effects caused spTMS and cTBS on the control of object manipulation (72).

For additional details and rationale for the control conditions, please refer to *SI Appendix*, S5.

### EEG data analysis

Dipole moments (pA*m) were used to quantify neural activity in the alpha band (9-12 Hz). Source maps were projected to a default brain (73) and subsequently decibel-normalized to a baseline period (average of −500 to −250 ms before a ‘ready’ cue) (Fig. 3*A*). After baseline normalization, data recorded between contact and object lift onset were averaged within subjects for each condition prior to statistical analysis. Source localization and analysis was performed using a combination of the Brainstorm toolbox (74) and Brainsuite (75) for cortical parcellation of individual subject structural T1-weighted MRIs (3T, Philips). The cortical surfaces for each subject were reconstructed from the MRIs using Brainsuite. MRIs were co-registered with EEG electrode locations, and used to create a boundary element model (BEM) of scalp, outer skull, and inner skull before source estimation. Source cortical activity from each trial was estimated using distributed source imaging based on a depth-weighted (L2 Norm) minimum-norm estimation (MNE) that estimated an orientation constrained dipole at each parcellated location from the BEM (total of 15,000 dipoles) (76) and converted to current density maps (77). Our subsequent analysis focused on two regions of interest (ROI): left precentral and postcentral gyrus, corresponding to M1 and S1, respectively. These ROIs were identified through automatic neuroanatomical labeling in Brainsuite (Fig. 3*A*).

### TMS data analysis

Electromyography (EMG) signals were recorded from the right FDI muscle using bipolar surface electrodes (Delsys Bagnoli system, Boston, MA) and digitized at 5 kHz (Power 1401 Cambridge Electronic Design, Cambridge, UK). Peak-to-peak MEP amplitudes (mV) were measured and extracted using a custom written Spike2 script and analyzed using Matlab. EMG signals were screened online and recorded during cTBS stimulation to verify that cTBS did not evoke MEPs.

### Behavioral data analysis

Data from force/torque sensors and the IMU gyroscope (range of ±2000º/s and noise density of 0.05 rad/s/√Hz) were sampled at 1000 Hz and 128 Hz, respectively. Force, torque, and object roll data were used to compute the following variables (Mathworks, Natick, MA): (1) *Digit forces*: Digit tangential force (F_tan_) is the vertical force component parallel to the grip surface produced by each digit to lift the object (Fig. 1*A,B*). Digit load force data exerted by each digit was used to compute the difference between thumb and index finger load forces (F_tan1_ − F_tan2_ = d_LF_). Digit normal force (F_n_) is the force component normal to the grip surface produced by each digit (Fig. 1*A,B*). Digit grip force was defined as the average of the thumb and index finger normal forces ([F_n1_ + F_n2_]/2 = F_GF_). (2) *Digit center of pressure*: The center of pressure of thumb and index fingertip (CoP_1_ and CoP_1_, respectively) was computed using the force and torque output of each sensor (25) (Fig. 1*A,B*). CoP data were then used to compute the vertical distance between the CoP on the thumb and finger side of the grip device (CoP_1_ – CoP_2_ = d_Y_). We computed the compensatory torque exerted on the object (T_com_, Fig. 1*A,B*) using the following equation:

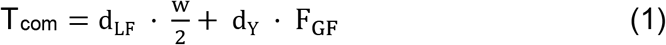

where ‘w’ denotes the width of the object. (3) *Peak object roll*: Our previous studies have demonstrated that T_com_ is a valid predictor of manipulation performance, i.e., object roll. Specifically, as subjects learn the appropriate T_com_ required to minimize object roll, peak object roll negatively correlates with the magnitude of T_com_ (25, 26, 31, 63). This was confirmed by a significant linear correlation between T_com_ and peak object roll (Pearson correlation coefficient on data pooled across all experimental groups and subjects: 0.68; *P* < 0.001). We also note that the results of the analysis of peak object roll and T_com_ were identical across all experimental and control groups. Therefore, as both variables capture two interrelated phenomena associated with learning dexterous manipulation, for the sake of brevity we report only results of the analysis of T_com_.

All of the above variables were computed at the time of object lift onset to quantify anticipatory control of manipulation (24, 25). Object lift onset was defined as the time at which the first of the two object switches was released from the object switch plate and remained open for 50 ms.

### Statistical analysis

#### EEG experiment

Baseline normalized dipole moments averaged within subjects for each grasping condition (30 trials each; Fig. 2*A*) were compared using t-tests with Bonferroni corrections. Our framework predicts that the neural drive from S1 to M1 should be greater for *uncon* compared to *con* grasping between contact and object lift onset. To this end, we used Granger causal (GC) analysis on the M1 and S1 source-reconstructed EEG data to determine the effective connectivity between these regions. The time epochs used to analyze these data were pulled from the same time window used to analyze source power. For each grasp context, the GC values between M1 and S1 were estimated using all trials for each subject individually. By analyzing all trials at the same time for a given subject and condition, we were able to determine whether there was an overall significant effective drive between M1 and S1 (and vice versa) in a subject-and condition-specific manner. GC was estimated using the Matlab MVGC toolbox (79). The toolbox fits multivariate vector autoregressive models up to a certain delay order. This model was estimated on a subject and condition basis to optimize the model fit. We detected significant GC in all subjects and conditions, all of which were used for analysis in a repeated measures ANOVA.

#### TMS experiment

We assessed subjects’ ability to perform the manipulation task by comparing T_com_ from the first trial with the average of the last five trials of each block (*Learn, Pre, Post*) within and across experimental groups (Fig. 2*B*). Our previous work has shown that subjects quickly learn to generate the necessary T_com_ (Fig. 4) within the first three trials so as to minimize object roll (25, 80). Analysis of the first trial of each block thus allowed the assessment of subjects’ performance without any previous experience (*learn1*), and recall of stored sensorimotor memory of grasp position and forces acquired after learning the manipulation task (*pre1*, *post1*). Subsequently, averaging the last five trials of each block (*learn5*, *pre5*, *post5*) was performed to obtain a measure of stability of performance for each block.

To assess learning-related changes in T_com_, we performed a 5 x 2 between-within repeated measures (rm) ANOVA with *Group* (5 levels: M1 *uncon*, M1 *con*, S1 *uncon*, S1 *con*, Vertex) as the between-subject factor, and *Block* (2 levels: l*earn1*, *learn5*) as the within-subject factor. To confirm that subjects’ performance remained stable during trials after learning and prior to cTBS, we performed a 5 x 3 between-within rmANOVA with *Group* as the between-subject factor, and *Block* (3 levels: *learn5*, *pre1*, *pre5*) as the within-subject factor. A similar statistical design was used to assess T_com_ in the remaining control groups (for details see S2 and S5, *SI Appendix)*.

To assess the effects of cTBS on T_com_, we performed a 5 x 3 between-within rmANOVA with *Group* as the between-subject factor, and *Block* (3 levels: *pre5*, *post1*, *post5*) as the within-subject factor. Post-hoc t-tests and one-way ANOVAs were used to compare between-and within-group differences, respectively. We performed separate one-way rmANOVA to assess the effects of cTBS on individual T_com_ components (d_LF_, d_y_, and F_GF_). We chose to perform only within-group analyses for individual position and force data because subjects could have selected different magnitude of the of T_com_ components giving rise to spurious between-group differences, even though the overall goal of T_com_ generation is the same. Finally, we used one-sample t-tests on the percentage changes (post-versus pre-cTBS, or following rest for the *No Move* group; *SI Appendix*, Fig. S1) in MEP data. We applied Huynh-Feldt corrections when sphericity assumption was violated. We used Dunnett’s post hoc test to compare each experimental group with the control group (i.e., Vertex). We used Bonferroni t-test for posthoc comparisons between experimental groups. For within-subject factors, we performed post hoc comparisons using paired t-tests with appropriate Bonferroni corrections.

Our analysis examining the difference in T_com_ of *pre5* and *post1* trials allowed us to quantify the immediate effect of cTBS to different neural sites. This revealed differential changes in T_com_ components (Fig. 5*A,B*). To understand whether these effects persisted or changed during the 15 post-cTBS trials, we calculated the difference of each post-cTBS trial and *pre5* data to create a time series of values. We notate these values with a ∆ to represent the difference; for example, the difference in d_Y_ pre-versus post-cTBS is denoted as ∆d_Y_ (*SI Appendix*, Fig. S3). To analyze the potential changes in these components as a function of *Group* over the time-course of *post-cTBS Trials* (1:15), Δd_Lf_, ΔF_GF_, and Δd_Y_ were analyzed using a repeated-measures mixed-model framework. *Group* entered the model as a between-subjects categorical factor, and *post-cTBS Trials* entered as a continuous covariate. Because we anticipated time-based effects due to the sequential nature of the task, we set the residual-covariance matrix to be scaled within subject and have an autoregressive (lag-1) structure. The model was also set to include random-intercepts for individual subjects. Each model was fitted using restricted maximum likelihood, and always started with a full structure with both the effects and interaction of *Group* and *post-cTBS Trial*. This full model was compared to each of the simpler, nested models of just the effects using a likelihood ratio test to determine the appropriate model. The final, reduced model is presented in S5, *SI Appendix*. All of the analyses were computed in the R environment using the lme4 (81) and lmerTest (82) packages.

## Contributions

All authors contributed to the design of the study, data analysis and interpretation, and manuscript preparation. PJP performed the TMS experiments, and JMF performed the EEG experiments. All authors approved the final version of the manuscript.

## Acknowledgments

We thank Dr. Patrick McGurrin for contributing to EEG and TMS data collection and analyses. This research was partially supported by a Collaborative Research Grant BCS-1455866 from the National Science Foundation (NSF) to MS and a grant from the Core for Advanced MRI (CAMRI) at Baylor College of Medicine to PJP. Its contents are solely the responsibility of the authors and do not necessarily represent the official views of NSF. We would like to thank Nishant Rao for assistance with data collection and MR technologist Lacey Berry, BS, RT(R)(MR) for assistance with MR scanning at the University of Houston. We thank Drs. Qiushi Fu, Jamie Tyler, Jeffrey Kleim, Andrew Gordon and Marco Davare for comments on the manuscript.

## Appendix: Supplementary Information

### S1. Rationale for design of dexterous manipulation task

Our dexterous manipulation task required subjects to learn to anticipate the effects of a destabilizing external torque (T_ext_) on the object by exerting a compensatory torque (T_com_) at object lift onset (1) (Fig. 1*A*). We chose to study task over the classic task of lifting an object with a symmetrical center of mass because the task goal of object roll minimization introduces an element of dexterity in addition to those that have been extensively studied, e.g., modulating normal force to load force to prevent object slip during lift and hold. Importantly, by combining our dexterity requirement with choice of contact points, we had earlier found that digit load force distribution is modulated to variable digit position prior to object lift onset on a trial-to-trial basis (see Introduction). It is important to note that this phenomenon of digit force-to-position modulation is also found when manipulating objects with a symmetrical mass distribution and there is no requirement for exerting a compensatory torque – in fact, in this scenario the covariation between digit load force distribution and position is even stronger than when manipulating objects with an asymmetrical mass distribution (see Fig. 8C (1)). These observations led to the proposition that digit force-to-position modulation is a task- and object-independent phenomenon underlying skilled manipulation through unconstrained grasp contacts (2).

### S2. TMS and EEG general procedures

The TMS coil was held tangential to the scalp, perpendicular to the direction of the central sulcus, 45° from the mid-sagittal line, with the handle pointing backward to induce current in the postero-anterior direction (3). Resting motor threshold (rMT) was defined as the TMS intensity that induced 50 μV peak-to-peak motor evoked potentials (MEPs) in 5 of 10 trials in the FDI muscle. Active motor threshold (aMT) was estimated by stimulating M1 at the same site used for rMT while the subject maintained a static contraction using the thumb and index finger on the object at approximately 20% of maximum voluntary contraction, defined as the average of three trials. We defined aMT as the TMS intensity that induced 200 μV peak-to-peak MEPs in 5 of 10 trials in the FDI muscle. For MEP analysis, we removed trials in which EMG activity during the 150-ms window prior to the single-pulse TMS was larger than 2 standard deviations of the mean baseline activity (calculated as the mean of the rectified EMG signal during a short period of rest). This was done to ensure that recorded MEP values were not affected by baseline EMG activity at the time of TMS stimulation (4).

Scalp EEG was recorded with a standard 10-20 layout. Sixty channels were recorded from the scalp with an AFz ground and left mastoid reference. Scalp electrodes FT9, FT10, PO9, and PO10 were used to record electrooculogram (EOG) signals, selected based on their distance from our regions of interest. EOG electrodes for horizontal eye movements were placed at the canthus of each eye, while vertical EOG electrodes were placed above and below the left eye. Trial epochs were then created using a time window (−1500 ms to 3000 ms) around the object contact event (time 0). Epochs containing large scalp EMG activity or where subjects did not comply with task instructions were excluded (< 10% of all trials). Electrodes showing abnormally noisy activity were interpolated using a spherical algorithm after applying independent components analysis for artifact rejection.

### S3. Digit force modulation to variable position occurs in unconstrained but not constrained grasping

During the *Learn* and *Pre*-cTBS trial blocks (Fig. 2*B*), subjects from all experimental groups learned to generate compensatory torque (Tcom) appropriate to minimize object roll. Learning of T_com_ occurred within the first three trials, after which T_com_ was consistently attained (Fig. 4*A*). Trial-to- trial modulation of digit load force distribution (d_LF_), grip force (F_GF_) and digit position (d_Y_) measured at lift onset was similar to that described in previous work (1, 5, 6). d_Y_ and d_LF_ from each trial were normalized to generate z-scores and used for linear regression analysis (1) to assess their relation in the EEG (*con* and *uncon* groups; Fig. 2*A*) and TMS experiments (M1 *con*, M1 *uncon*, S1 *uncon*, S1 *con*, and Vertex groups; Fig. 2*B*). Z-scores were computed by normalizing d_LF_ and d_Y_ for each subject by removing the mean from the value of each trial and dividing the result by the standard deviation.

As T_com_ is learned within the first three trials (1, 5), we used all *con* and *uncon* trials after learning had occurred for the EEG experiment (30 trials for each grasp context per subject), and trials 4-10 of the *Learn* block and all trials in the *Pre* block (22 trials per subject) for the TMS experiment. As expected from our previous work, we found (a) higher d_y_ variability in all *uncon* than *con* grasping conditions from the EEG and TMS experiments (all *P* < 0.05), and (b) the larger d_Y_ variability in *uncon* was compensated by trial-to-trial modulation of d_LF_ (25, 27, 30). Specifically, we found significant negative correlations between d_LF_ and d_Y_ only for the *uncon* grasping condition. For the EEG experiment, the *r-*value was −0.51 (*P* < 0.001) for *uncon* and −0.096 (*P* = 0.104) for *con*. For the TMS experiment, we found significant negative correlations between d_LF_ and d_y_ in M1 *uncon*, S1 *uncon*, and *Vertex* conditions (*r* = –0.45, –0.67, and –0.46; all *P* < 0.0001), but not in M1 *con* and S1 *con* groups (*r* = 0.08 and −0.12, respectively; all *P* > 0.1).

### S4. cTBS to M1 and S1 does not reduce corticospinal excitability following exposure to object manipulation

We found no change in corticospinal excitability (CSE) after continuous theta burst stimulation (cTBS) was delivered over M1 and S1 in the experimental groups (M1 *con*: t_9_ = −2.052, M1 *uncon*: t_9_ = –2.314, S1 *con*: t_9_ = −0.98, and S1 *uncon*: t_9_ = −0.991, respectively; all *P* > 0.05), nor M1 in the control groups (*Sham*, *no-cTBS*, and *Vertex*: all *P* > 0.05; Fig. S1).

These findings may seem surprising, as previous work reported a reduction in CSE following cTBS to M1 (7). Unlike our protocol, however, in this previous work subjects did not perform a motor task prior to M1 cTBS. This is an important methodological difference, as a later study by the same group reported no reduction in CSE when subjects performed an isometric force contraction during cTBS stimulation (8). Therefore, the lack of CSE reduction following cTBS in our study, where subjects performed a series of object lifts prior to cTBS, is consistent with the follow-up study by Huang and colleagues (8). Nevertheless, to further validate our cTBS protocol, we performed an additional test on a *No move* group (*n* = 6) where we assessed the effects of cTBS over M1 on MEP size without having subjects perform our manipulation task (Fig. 2*B*). In this group and consistent with previous work where subjects performed no motor tasks prior to M1 cTBS (7), we found a significant decrease in MEP amplitude (t_9_ = −7.172, *P* = 0.001; Fig. S1).

### S5. cTBS delivered to control groups does not affect compensatory torque

In addition to the *Vertex* group, we ran three additional control experiments. In each of the control groups described below, participants were given the same task instructions given to subjects in the four experimental groups (Fig. 2*B*).

#### Sham (n = 10)

CSE was assessed using single-pulse TMS (spTMS) over contralateral (left) M1 region immediately after the *Pre* block and before the start of the *Post* block, corresponding to the time immediately before and 5 minutes after cTBS. cTBS was delivered to a second coil placed immediately behind subject’s head with current directed away from the scalp while the coil over contralateral (left) M1 remained in place. Thus, subjects heard the sound elicited by stimulation, but did not experience any somatosensory effect of stimulation on the scalp. This control group was used to control for any somatosensory effects (9) caused by the auditory cue of cTBS on the control of object manipulation.

#### No cTBS (n = 6)

This group did not receive cTBS. CSE was assessed using spTMS over contralateral (left) M1 region immediately after the *Pre* block and before the start of the *Post* block. This was done to assess the influence of MEP-induced movements on object manipulation control. Muscle twitches caused by spTMS over M1 have been shown to affect grasping behavior in subsequent lifts (10). Therefore, the results of this control condition were analyzed to ensure that any change in behavior found in the experimental groups was specifically due to a ‘virtual lesion’ over the cortical area targeted by cTBS.

#### No Stim (n = 6)

Neither spTMS nor cTBS were delivered during the experiment. To assess learning-related changes in T_com_ in control groups, we performed a 4 x 2 between-within repeated measures (rm) ANOVA with *Group* (4 levels: *Sham, no cTBS, No Stim, Vertex*) as the between-subject factor, and *Block* (2 levels: *learn1*, *learn5*) as the within-subject factor. We chose to include the *Vertex* group in this analysis to ensure that there were no differences across any of the control groups. This inclusion also served to validate that having included any control groups in the main analysis with the M1 c*on*, M1 *uncon*, S1 *con* and S1 *uncon* groups would have produced similar results. We report a significant main effect of *Block* (*F*_1,28_ = 320.46, *P* < 0.05), but no significant *Block × Group* interaction (*F*_1,28_ = 2.50, *P* > 0.05). Individual rmANOVAs for each group confirmed that the magnitude of T_com_ significantly increased by the end of the *Learn* block (learn1 versus learn5, Fig. S2) for all control groups (significant main effect of *Block*, all *P* < 0.05).

In addition, we confirmed that object manipulation performance remained stable during trials after learning and prior to cTBS using a 4 *×* 3 between-within rmANOVA with *Group* (4 levels: *Sham, no-cTBS, No Stim, Vertex*) as the between-subject factor, and *Block* (3 levels: *learn5*, *pre1*, *pre5*) as the within-subject factor. After learning, T_com_ remained invariant at the beginning of the *Pre* block (no main effect of *Block*: *F*_2,56_ = 0.267, *P* > 0.05; no significant *Block × Group* interaction: F_2,56_ = 0.572, *P* > 0.05; Fig. S2). This comparison confirmed both that the rest period between the *Learn* and *Pre* blocks had no significant effect on the learned T_com_ (learn5 vs. pre1) and that T_com_ was stable throughout the *Pre* block (pre1 vs. pre5) (Fig. 4*B*). These results are identical to those reported for the experimental groups.

To identify differences across trial blocks and control groups, we used a 4 *×* 3 between-within rmANOVA with *Group* (4 levels: *Sham, no-cTBS, No Stim, Vertex*) as a between-subject factor and *Block* (3 levels: pre5, post1, post5) as the within-subject factor. We found no main effect of *Block* (*F*_2,56_ = 2.73, *P* > 0.05) nor significant *Block × Group* interaction (*F*_2,56_ = 0.42, *P* > 0.05). Lastly, between-group comparisons for all control groups revealed no differences during the *Learn, Pre, or Post* block trials (all *P* > 0.05, Fig. S2). Therefore, subjects in all control groups attained and maintained similar T_com_ throughout the remainder of the *Pre* and *Post* blocks.

Together, these analyses confirms that cTBS to sites other than M1 and S1, and/or the presence of spTMS between blocks, did not affect skilled object manipulation performance. Similar findings were found for peak object roll (see S5).

### S6. Persistence of cTBS effects on compensatory torque components is sensitive to grasp context and cortical area

Analysis of the first post-cTBS trial, as well as subsequenty post-cTBS trials, revealed a differential effect on T_com_ components depending on the cortical area being stimulated and grasp context (Fig. 5*A*,*B*). Therefore, we quantified how these changes in T_com_ components persisted over post-cTBS trials. We used mixed models to examine changes in each T_com_ component over post-cTBS trials using a factor of Group (M1 *con*, M1 *uncon*, S1 *uncon* and S1 *con*) and a continuous covariate of post-cTBS Trial (1-15). For each T_com_ variable, the model used the difference (Δ) of the average of pre5 trials and each post-cTBS trial as the dependent variable.

The results for Δd_LF_ revealed a significant interaction between Group and post-cTBS Trial (F_1,3_ = 3.25, *P* < 0.05). The interaction resulted from a significant difference in slopes between M1 *uncon* and S1 *uncon* groups (*P* < 0.05; Fig. S3, top row). Follow-up examination revealed that both M1 *uncon* and S1 *unco*n exhibited slopes that were different from 0 (both *P* < 0.05). In contrast, the model examining ΔF_GF_ only revealed an effect of Group (F_1,3_ = 8.67, *P* < 0.05). This Group effect is apparent in Fig. S3 (middle row), as the mean level of the line for each group differs at a steady-state across post-cTBS trials. More specifically, follow-up analysis of the intercept revealed only the M1 *con* group differed from zero (*P* < 0.05). Therefore, M1 *con* was the only experimental condition with the strongest and more persistent effects of cTBS on F_GF_. The final model examining Δd_y_ also yielded a significant interaction (F_1,3_ = 6.95, P < 0.05; Fig. S3, bottom row). Follow-up analysis revealed that only the slope for M1 *uncon* and S1 *con* groups were significantly different from 0, with both exhibiting a positive trend post-cTBS (both *P* < 0.05).

These findings indicate that the effects of cTBS on digit forces and positions were highly sensitive to the grasp context and cortical area targeted by TMS. Specifically, the persistency of the effects of virtual lesion on T_com_ for M1 *con* grasping throughout all post-cTBS trials (Fig. 5*B*) can be solely attributed to alteration of F_GF_, as indicated by the persistent and large non-zero intercept across post-cTBS trials (M1 *con*, Fig. S3). In contrast, the faster recovery of T_com_ to pre-cTBS levels for M1 *uncon* grasping can be attributed to the re-establishment of a negative covariation between d_y_ and d_LF_ (M1 *uncon*, Fig. S3), despite large trial-to-trial fluctuations in d_y_ (Fig. 5*B*). For the S1 *con* group, the quick recovery of T_com_ to pre-cTBS level after the first post-cTBS trial was mediated only by adjustments in relative positioning of thumb and index finger within the marked contact boundaries on the object (Fig. S3). Lastly, the rate at which T_com_ recovered within the first 10 post-cTBS trials (Fig. 5*B*) following cTBS in the S1 *uncon* group was mostly driven by change in d_LF_ (right column, Fig. S3).

---

**Fig. S1.**
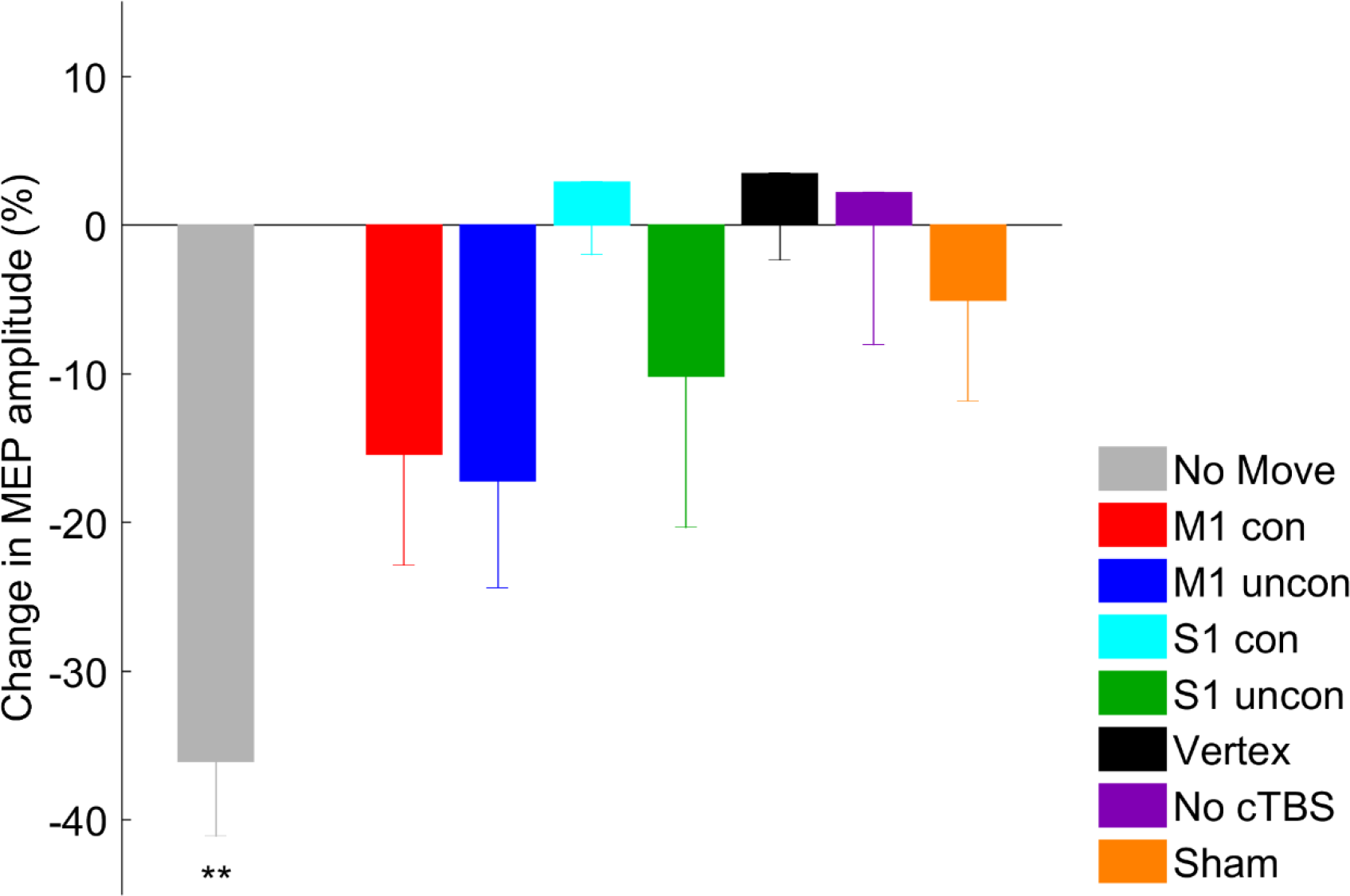
Corticospinal excitability. Change in CSE was assessed as percentage change in the amplitude of motor evoked potentials (MEP) by comparing pre-versus post-cTBS, or following rest (No Move group). All groups except the No cTBS group received cTBS over M1, S1, or Vertex. ** denotes *P* < 0.0125. Data are averages (± SE) of all subjects.

**Fig. S2.**
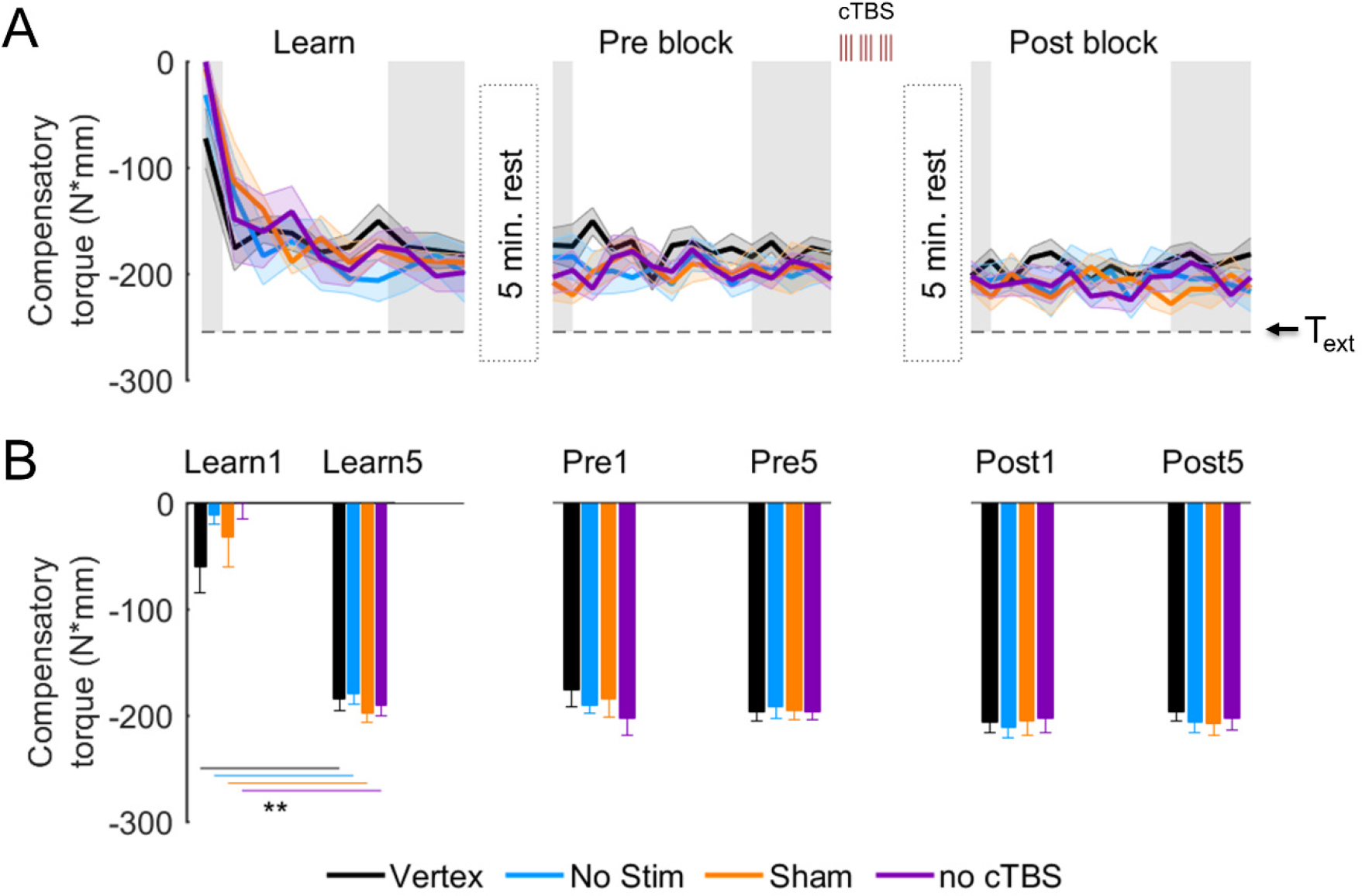
Compensatory torque: Control groups. (*A*) Compensatory torque (T_com_) during Learn, Pre, and Post blocks in the Vertex, No Stim, Sham, and No cTBS groups. (*B*) T_com_ on the first trial and the average of the last 5 trials for each block. Data are plotted in the same format as Figure 4. ** denotes *P* < 0.0125.

**Fig. S3.**
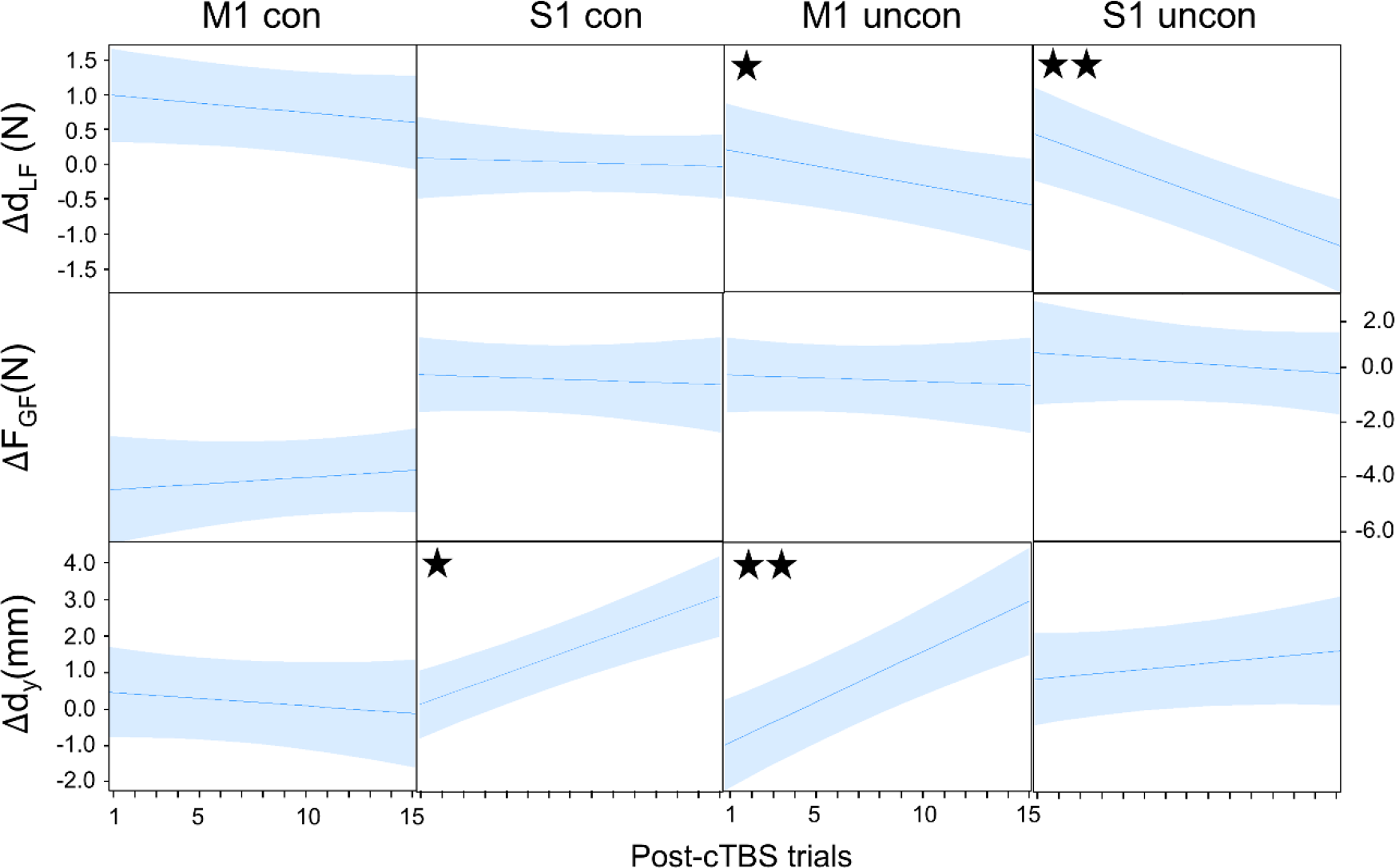
Effect of cTBS on digit placement, load and grip force. Plots show predicted difference (Δ) between the value of each T_com_ variable averaged across the last five pre-cTBS trials and each post-cTBS trial. Predicted values were obtained by fitting a mixed model that predicted the variable (e.g., d_LF_) as a function of experimental group and post-cTBS trial. Each plot shows the predicted slope. ✭ and ✭✭ denote a slope significantly different than zero at *P* < 0.05 and 0.01, respectively.

